# Intercellular Communication via Mitotic Nanotubes is Influenced by Connexin-43 Trafficking and Actin Remodeling

**DOI:** 10.64898/2026.02.08.704470

**Authors:** A. Cooper, S. Cetin-Ferra, K.A. Yonosh, A. Hinton, A. Marshall, J. R. Faeder, S.A. Murray

## Abstract

Gap junction communication is reduced during mitosis as the junction protein connexin-43 (Cx43) is redistributed from gap junction plaques on the plasma membrane to cytoplasmic annular vesicles and actin-based mitotic nanotubes that transiently connect mitotic cells to neighboring cells. However, the dynamic details of Cx43 redistribution during cell entry into and exit from mitosis, and the roles of mitotic nanotubes and associated Cx43 in intercellular communication, remain poorly understood. Here, using confocal live-cell imaging, we show that as cells enter mitosis, plaque-derived Cx43 structures are transferred to mitotic nanotubes. Over time, these structures fragment and migrate along the length of the nanotubes, either being transferred to the cytoplasm of adjacent cells or being positioned at the nanotube ends where they could potentially enable communication. Functionally, mitotic nanotubes indeed facilitate gap junction-dependent intercellular communication, though at reduced rates compared interphase cells. Interestingly, knockdown of Cx43 resulted in impaired nanotube formation and intercellular communication while inhibition of Rho kinase (ROCK) with Y-27632 prevented mitotic cell rounding and nanotube elongation, and increased cell–cell communication during mitosis, suggesting that nanotube function is influenced by Cx43 expression and trafficking as well as actin remodeling via ROCK. Overall, these findings provide valuable insights into the mechanisms that regulate Cx43 and mitotic nanotube dynamics and reveal a novel role for mitotic nanotubes in facilitating cell–cell communication during cell division.

## Introduction

Gap junction channels are protein complexes that span the plasma membranes of apposing cells, thus allowing for the direct passage of small molecules and ions. The formation of these channels is a well-defined process that involves: (1) synthesis of the gap junction channel proteins, connexins; (2) assembly of connexins into hemichannels on the plasma membranes; and (3) docking of the hemichannels from apposing cells to form gap junction channels at the interfaces between adjacent cells [1–7]. While there are 21 connexin proteins expressed in human tissues, connexin 43 (Cx43) is the most widely expressed and extensively studied gap junction protein [8]. Following docking, gap junction channels aggregate to form gap junction plaques that allow for more efficient cell–cell communication than single channels [4, 7, 9]. In addition to channel gating (opening and closing), gap junction-mediated intercellular communication is largely regulated by the removal of gap junction plaques from the plasma membrane [7, 10, 11]. The canonical pathway for this process involves clathrin-dependent internalization of gap junction plaques, resulting in the release of double-membrane vesicles, called annular gap junctions, into the cytoplasm of either of the two adjacent cells [7, 12–14]. Annular gap junctions have been shown to fuse with lysosomes, which leads to the degradation of the connexin proteins [12, 15] and is therefore considered to be a key step in the gap junction protein turnover pathway.

While gap junction turnover has been extensively characterized in cell populations that primarily contain interphase cells, our understanding of these processes during mitosis is limited. However, there is a prevailing view that intercellular communication via gap junctions is reduced compared to that in interphase cells [16–18]. The extent to which intercellular communication is reduced as a function of gap junction distribution changes throughout the mitotic cycle remains unclear. Moreover, although gap junction-mediated communication may be diminished during mitosis, the functional significance and mechanisms of any residual communication remain unknown. We and others have reported a decrease in gap junction plaques accompanied by an increase in annular gap junctions, changes that were speculated to contribute to the mitosis-associated reduction in gap junction-mediated communication [19–21]. The possibility that a small amount of communication persists during mitosis, and the potential benefits of this residual communication for cell behavior, remain unexplored. Together, these observations raise important questions about the fate of gap junctions during and after mitosis, and particularly how gap junction communication is reestablished following cell division. It has been speculated that, after mitosis, annular gap junctions recycle to the plasma membrane to form new gap junction plaques as cell–cell communication is reestablished in the new daughter cells [19–21].

Fykerud and colleagues demonstrated that during mitotic cell rounding, actin-based plasma membrane bridges formed between mitotic and adjacent cell pairs [20]. These extensions, coined “mitotic nanotubes,” contained both Cx43-positive vesicles and Rab11-positive recycling endosomes, thus leading to the speculation that Cx43-positive vesicles are transiently stored in mitotic nanotubes and then delivered to the plasma membrane to form gap junction plaques in newly formed daughter cells [20]. However, given that previous reports are based on imaging fixed cells or live cells with limited vesicle tracking [20], further analysis is needed to demonstrate this process definitively. In addition, studies are needed to further characterize these mitotic nanotubes by pinpointing the cell cycle stages at which they are formed and lost, and to assess dynamic changes in their morphology and their associations with Cx43-structures. Importantly, the role of mitotic nanotubes in gap junction-dependent communication between mitotic cells and their neighbors needs to be identified. Therefore, while recent findings in the field have increased our understanding of mitosis-associated changes in gap junction intercellular communication, critical gaps in this area remain, and further investigations are needed to determine the role of mitotic nanotubes in Cx43 trafficking and cell–cell communication, thereby improving our understanding of gap junction function during mitosis.

Specifically, real-time analysis of gap junction proteins, as cells both enter and exit mitosis, is necessary to further our understanding of Cx43 fate and function in intercellular communication throughout the cell cycle. In addition, while changes in Cx43 distribution have been reported in multiple cell types during mitosis [19–21], these demonstrations lack the quantification required for statistical analyses and for developing mathematical models of gap junction protein behavior throughout the cell cycle. Beyond our understanding of Cx43 localization to gap junction plaques and annular gap junctions, the redistribution of Cx43 into mitotic nanotubes is a recent report and therefore largely unexplored, having not been quantified in previous assessments of Cx43 subcellular distribution. Yet such information is needed to further assess the formation, function, morphology, and composition of these structures over time.

In this study, three-dimensional confocal time-lapse imaging was performed to visualize gap junctions in an adrenal-derived cell line transiently transfected to express fluorescent-tagged Cx43 (mEmerald-Cx43) throughout the cell cycle. We demonstrate the mechanism of mitotic nanotube formation and connexin association during early mitotic stages and quantify Cx43 redistribution as cells progress from interphase to cytokinesis. Importantly, we identify a novel role for Cx43-associated mitotic nanotubes in maintaining gap-junction-mediated intercellular communication and establish how Cx43 trafficking and actin remodeling influence this role. Thus, these findings provide valuable insights into the mechanisms that influence mitosis-associated changes in intercellular communication and establish mitotic nanotubes as potential targets for specifically influencing mitotic cells.

## Methods

### Cell Culture

The human adenocarcinoma cell line, SW-13, was obtained from ATCC and grown as previously described [10, 21]. Briefly, cells were cultured in Lebovitz L-15 media (Gibco #41300-021) supplemented with 10% fetal bovine serum, 200 U/ml penicillin, 200µg/ml streptomycin, and 5mg/ml Amphotericin B (Fungizone). Cells were maintained at 37°C in an incubator without carbon dioxide and passaged at 1:5 to 1:20 dilution to maintain culture. For experimental analysis, cells were used at or below passage 16.

### Treatment with Inhibitors

Gap junctions were inhibited by treating cells with 3mM Octanol for 10 minutes. To inhibit ROCK, cells were treated with 10µg/ml Y-27632 (Sigma Aldrich, St. Louis, MA, #688000) for 2h. Control samples for cells treated with Y-27632 were treated with an equivalent concentration of the diluent DMSO.

### Cell Synchronization

Cells were synchronized using a double-thymidine block that was modified for SW-13 cells based on previously published protocols [22, 23]. Specifically, cells were seeded and allowed to adhere to the substratum before treatment. To perform the synchronization, cells were treated with 2 mM thymidine for 36 h (first thymidine block), then washed three times with cell culture medium to remove thymidine and incubated for 16 h (first release). Next, cells were incubated for a second time with 2mM thymidine for 36h (second thymidine block) and washed with cell culture medium to remove thymidine (second release). Cells were typically observed to undergo mitosis 12-16h following the release.

### Transfection

To visualize gap junctions with time-lapse microscopy, cells were transiently transfected by mixing Lipofectamine 2000 (Invitrogen, Carlsbad, CA, #11668030) and plasmid DNA in a 1:2 ratio in OPTI-MEM transfection medium (ThermoFisher, Waltham, MA, # 31985070) and applying this mixture to cells that were seeded on poly-lysine-coated glass-bottom dishes. Plasmids encoding the gap junction markers mEmerald-Cx43 (Addgene, Watertown, MA, plasmid #54055) and Halo-Cx43 were gifts from Michael Davidson and John O’Brien, respectively. The plasmid encoding the membrane marker mCherry-CAAX was a gift from Gerry Hammond. After 4 hours of incubation at 37 °C, the Lipofectamine-DNA mixture was removed and replaced with complete medium without antibiotics. For knockdown experiments, cells were transfected with either control (non-targeting) siRNA (IDT, Coralville, IA, #51-01-14-04) or siRNA targeted to human GJA1 (Cx43) gene (IDT, Coralville, IA, #hs.Ri.GJA1.13.1). Cells in 24-well plate format were transfected using 1ul Lipofectamine and 40nM siRNA. For cells synchronized with a double-thymidine block, transfection was performed during the first thymidine release.

### Western Blotting

Cells were lysed with cell extraction buffer containing protease inhibitor cocktail (Life Technologies, Carlsbad, CA) and PMSF and lysates were incubated with Laemmli buffer and ran on 15% SDS-PAGE gels, transferred to PVDF membranes, and probed with rabbit polyclonal anti-Cx43 primary antibody (Millipore, Burlington, MA, #AB1728) and HRP-conjugated anti-rabbit IgG secondary antibody (Jackson Laboratories, West Grove, PA, #111-035-003). Western blots were incubated in Chemiluminescent HRP substrate (MilliporeSigma, Burlington, MA, #WBKLS0100) and detected with a ChemiDoc imaging system with Image Lab Software Version 2.3.0.07 (BioRad Hercules, CA). Experiments were performed in triplicate, and representative Western blot images are presented.

### Immunocytochemistry

Immunocytochemistry was performed as previously described [21, 24]. Cells were fixed with 4% paraformaldehyde, permeabilized with acetone, and blocked with PBS containing 10% normal goat serum and 1% bovine serum albumin. Cells were stained with primary antibodies for 1 hour at room temperature, then with secondary antibodies and, in some cases, with phalloidin for 30 minutes at room temperature, and finally with DAPI nuclear stain. Cells were washed 3 or more times with PBS between each step. The coverslips were mounted onto glass slides using Fluormount G (Invitrogen, Carlsbad, CA, #00-4958-02). Antibodies and stains are listed in Table 1.

**Table 1.** Reagents used for immunocytochemistry.

| Reagent | Target | Supplier | Catalog Number | Dilution |
| --- | --- | --- | --- | --- |
| rabbit anti-Cx43 antibody | Cx43 | Sigma-Aldrich, Burlington, MA | AB1728 | 1:100 |
| mouse anti-Cx43 antibody | Cx43 | Sigma-Aldrich, Burlington, MA | MAB3067 | 1:100 |
| Alexa Fluor 488-goat anti-mouse IgG antibody | Mouse IgG | Invitrogen, Carlsbad, CA | A11029 | 1:500 |
| Alexa Fluor 488-goat anti-rabbit IgG antibody | Rabbit IgG | Invitrogen, Carlsbad, CA | A11034 | 1:500 |
| Alexa Fluor 647-goat anti-rabbit IgG antibody | Rabbit IgG | Jackson Immunology | 111-605-003 | 1:250 |
| Cy5-goat anti-mouse IgG antibody | Mouse IgG | Invitrogen, Carlsbad, CA | A10524 | 1:250 |
| Alexa Fluor 594-phalloidin | F-actin | Invitrogen, Carlsbad, CA | A12381 | 1:50 |
| Alexa Fluor 488-phalloidin | F-actin | Invitrogen, Carlsbad, CA | A12379 | 1:50 |
| DAPI | DNA | Invitrogen, Carlsbad, CA | D1306 | 1:5000 |

### Confocal Microscopy of Live and Fixed Cells

For imaging studies, confocal z-stack images were acquired with a Nikon A1 or Nikon AX confocal microscope with a 60x oil objective. Nikon NIS Elements software was used for image acquisition. For time-lapse microscopy of live cells, thirty z-slices 1µm apart were acquired at 5-minute intervals unless otherwise noted. Time is shown in the format hh: mm or in minutes. Images are presented as maximum intensity projections of multiple z-slices unless otherwise noted.

### Transmission Electron Microscopy

Cells were cultured and prepared for TEM images as previously described [10]. Briefly, cells were fixed in PBS containing 2.5% glutaraldehyde, then post-fixed in a solution of 1% osmium tetroxide and 1% potassium ferricyanide for 1 hour at 4 °C. Cell samples were embedded in resin, sectioned, mounted, and imaged with a JEOL 1011CX electron microscope.

### Analysis of Time-lapse Images

Maximum-intensity Z-projections were prepared and denoised using NIS Elements. For cell segmentation, Python was used to iterate over each time series and generate cell masks using Cellpose. Cell masks were manually corrected in Fiji ImageJ as needed. From each cell mask, the perimeter was used to generate plasma membrane masks, while the interior was used to generate cytoplasm masks. The plasma membrane areas between each pair of daughter cells and the mitotic nanotubes were manually segmented in ImageJ. Gap junction structures were segmented by automatic thresholding and objects <0.01µm^2^ were excluded to avoid improper analysis of noise or Cx43-containing secretory vesicles, shown to be smaller than annular gap junctions [7, 25, 26]. The morphometric features of gap junction structures in each cell at each time point were measured programmatically using Python.

### Fluorescence Recovery After Photobleaching (FRAP)-based Intercellular Communication Assay

A FRAP assay, adopted from methods previously used in the lab [24], was performed to assess the rate of intercellular communication. To assess the intercellular communication of calcein, cells were seeded on glass-bottom dishes and allowed to adhere overnight. The following day, cells were loaded with dye by incubating with 2µM calcein AM (Invitrogen, Carlsbad, CA #C3100MP) for 15 minutes at 37°C, then washed five times with serum-free media to remove calcein from the cell culture media. To measure intercellular communication of enhanced green fluorescent protein (EGFP), cells were transfected to express EGFP. The following day, transfected cells were seeded onto glass-bottom dishes and allowed to adhere overnight before performing the FRAP assay.

For cells loaded with calcein or EGFP, the FRAP assay was performed using a Nikon A1 confocal microscope with a 60x objective lens. For each field of view, an entire cell was selected as a region of interest (ROI) and photobleached with a 488nm laser at 100% laser power for 20-30s. Following bleaching, cells with calcein were imaged at 10s intervals for 150s, and cells expressing EGFP were imaged at 4s intervals for 60 seconds using the 488nm laser line at ≤2% laser power. Cells assessed with FRAP were in either of the three following categories: (1) isolated interphase cells (negative controls for gap junction communication), (2) interphase cells in contact with other interphase cells (positive controls for gap junction communication) or (3) mitotic cells that only associated with interphase cells via mitotic nanotubes. For FRAP with calcein-loaded cells, a minimum of 6 cells were assessed for each cell category; for FRAP with EGFP-expressing cells, a minimum of 3 cells were assessed for each cell category.

For each replicate, fluorescence from a background ROI was subtracted from the bleached cell ROI. The background-corrected values for the bleached ROI were divided by a background-corrected reference ROI to correct for changes in fluorescence due to photobleaching or drift. Data is expressed as percent fluorescence recovery compared to the pre-bleach values.

### Statistical Analysis

GraphPad Prism 10 software was used to generate graphs and perform statistical analysis. All data is presented as mean values. All error bars represent the standard error of the mean (S.E.M.). For comparisons between two groups, a student’s t-test was used to determine statistical significance; for comparisons between two or more groups, ANOVA with Tukey’s post-hoc multiple comparisons test was used. For graphs shown, asterisks (*) represent the appropriate p-values: 0.12 (ns), 0.033 (*), 0.002 (**), <0.001(***).

## Results

### Characterization and Distribution of Gap Junction Structures in Adrenal Cells Across the Cell Cycle

In interphase cells, gap junction plaques, and annular gap junction vesicles were detected and distinguished from one another with several microscopic techniques, including confocal immunocytochemistry to visualize endogenous Cx43, confocal live-cell imaging of cells transfected to express fluorescently tagged Cx43, and TEM (Figure 1A-E). With light microscopy, morphology, distribution, and abundance of gap junction plaques and annular gap junctions were observed, whereas TEM allowed appreciation of their ultrastructural details. Specifically, the characteristic double membranes of both gap junction plaques and annular gap junction vesicles, separated by a ∼2 nm gap, were clearly observed, along with the central lumen of the annular gap junction vesicles (Figure 1D, E). Such morphology is consistent with the annular gap junction vesicle membrane being derived from the internalization of a gap junction plaque.

**Figure 1.**
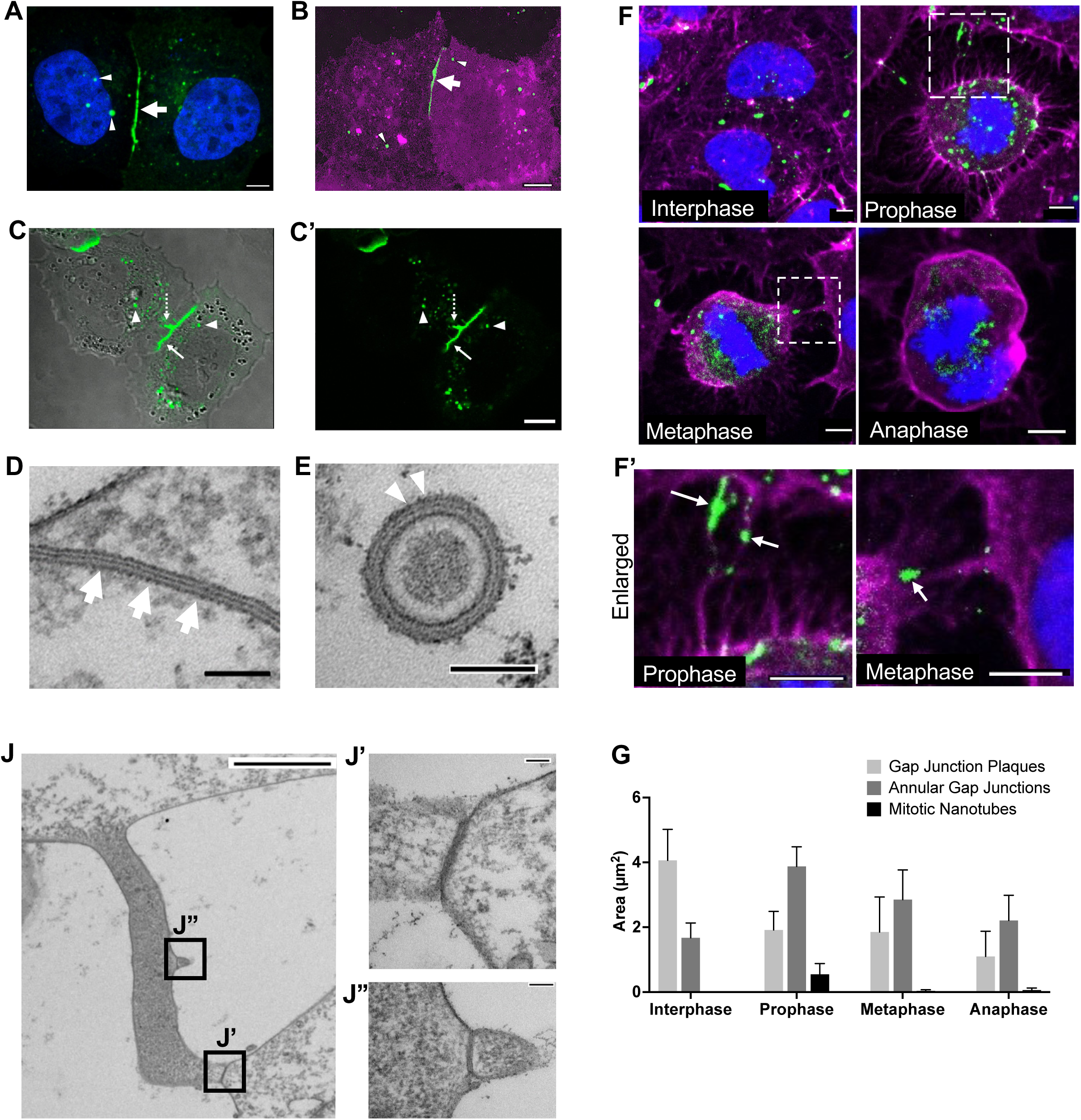
Characterization and distribution of gap junction structures in SW-13 cells across the cell cycle. (A) Confocal immunocytochemical image of fixed cells stained to visualize endogenous Cx43 (green) and nuclei (blue). (B) Confocal live-cell image of cells transfected to express mEmerald-Cx43 (green) and the plasma membrane marker mCherry-CAAX (magenta). (C and C’) Confocal live image of cells transfected to express Halo-Cx43 (green), shown with (C) or without (C’) DIC overlay. (D and E) Transmission electron micrographs of a gap junction plaque (D) and an annular gap junction (E) in which the typical pentalaminar membrane morphology can be distinguished. In A-E, solid arrows represent gap junction plaques; dashed arrows represent the invaginated region of the gap junction plaque; arrowheads represent annular gap junctions. (F) Cells in various cell cycle stages were fixed and stained to visualize actin (magenta), Cx43 (green), and nuclei (blue). (F’) Enlargements of mitotic nanotubes in outlined areas in F to show association with Cx43-positive structures (arrows). (G) Cellular distribution of Cx43-positive structures across cell cycle stages. The bar graph represents the mean total area of Cx43 structures per compartment per cell ± SEM. Across 3 experiments, 79 cells were analyzed, with n ≥ 8 per cell cycle stage. (J) Micrograph of a membrane extension that forms a bridge between two cells. The extension is closed at the distal end (J’) where it makes contact with another cell. Contact sites shown in A (boxes J’ and J’’) were magnified to show the typical gap junction plaque membrane. (J’) Contact site between the closed end of the membrane extension and the adjacent cell. (J’’) Contact site between an area along the length of the membrane extension and the tip of another membrane extension. Scale bars = 5µm (A, F, F’), 10µm (B, C, C’), 100nm (D, E), 2µm (J), 100nm (J’, J’’).

To assess gap junction structure distribution across the cell cycle, SW-13 cells were fixed and stained to visualize Cx43 at various cell cycle stages. Although Cx43 was predominantly localized to gap junction plaques between interphase cells, Cx43 localized mainly to annular gap junctions in cells in prophase, metaphase, and anaphase (Figure 1F, G). During mitosis, Cx43-positive structures were also observed to associate with actin-based mitotic nanotubes that formed bridges between mitotic cells and their neighbors (Figure 1F). The association of Cx43 and mitotic nanotubes was most prominent during prophase but was also observed to a smaller degree during metaphase and anaphase. Connexin-43 was detected in various linear and rounder, blob-like structures along the length of the mitotic nanotubes (Figure 1F, F’). Differences in the abundance and morphology of mitotic nanotube-associated Cx43 structures across mitotic stages are suggestive of dynamic remodeling of these gap junction structures.

To further assess mitotic nanotube morphology, cells were visualized with TEM. A membrane extension that is speculated to be a mitotic nanotube was observed in an unsynchronized SW-13 cell population (Figure 1J). The typical gap junction plaque morphology (pentalaminar structure) was observed at one end of the membrane extension (Figure 1J, J’). This morphological localization of the gap junction relative to the extension raises the hypothesis that during mitosis, at least one end of the tube could allow for gap junction-mediated intercellular communication. In contrast, the other end of the tube appeared to be open and continuous with the cell body of the cell it protruded from. At one point along the length of the extension, a gap junction plaque was observed where the extension contacted another smaller projection (Figure 1J, J’’), thus suggesting that mitotic nanotubes may form direct interactions and points of intercellular transfer with other mitotic nanotubes. Given the limited field of view in TEM, it is not possible to determine the cell cycle stage of the cells at either end of the membrane extension; therefore, it cannot be confirmed whether this extension meets the definition of a mitotic nanotube. However, these observations, along with immunocytochemistry studies, provide insight into the structural features of mitotic nanotubes and support the possibility that these structures contain gap junctions that may facilitate communication despite reductions in gap junction plaques during mitosis. Altogether, these data demonstrate a relationship between cell cycle stage and Cx43 distribution and establish mitotic nanotubes as a reservoir for Cx43 during mitosis. However, further investigations are needed to elucidate the dynamics of Cx43 redistribution to mitotic nanotubes.

### Dynamic Redistribution of Gap Junction Structures During Mitosis

To visualize gap junction trafficking, cells expressing mEmerald-Cx43 and the membrane marker mCherry-CAAX were synchronized and imaged for >16h at 5-minute intervals. We sought to assess gap junction turnover in interphase cells as they prepared for mitosis. In interphase cells that did not divide throughout the imaging period, portions of gap junction plaques were observed to internalize into either of the two contacting cells to form annular gap junctions (Figure 2A). Still, the internalization process did not appear to significantly alter the size of gap junction plaques on the surface between cell pairs (Figure 2A). The stability of gap junction plaque amounts at the cell surface is consistent with a dynamic balance between plaque downregulation via internalization and degradation and plaque formation via new Cx43 synthesis and assembly. However, in interphase cells that later entered mitosis, gap junction plaque internalization occurred more frequently, resulting in a larger number of annular gap junctions being released into the cytoplasm compared to that in non-dividing cell pairs (Figure 2A). Thus, while gap junction plaque levels in non-dividing cells remained stable over several hours, cells preparing to divide showed increased gap junction plaque internalization, resulting in progressive decreases in gap junction plaques at the cell surface.

**Figure 2.**
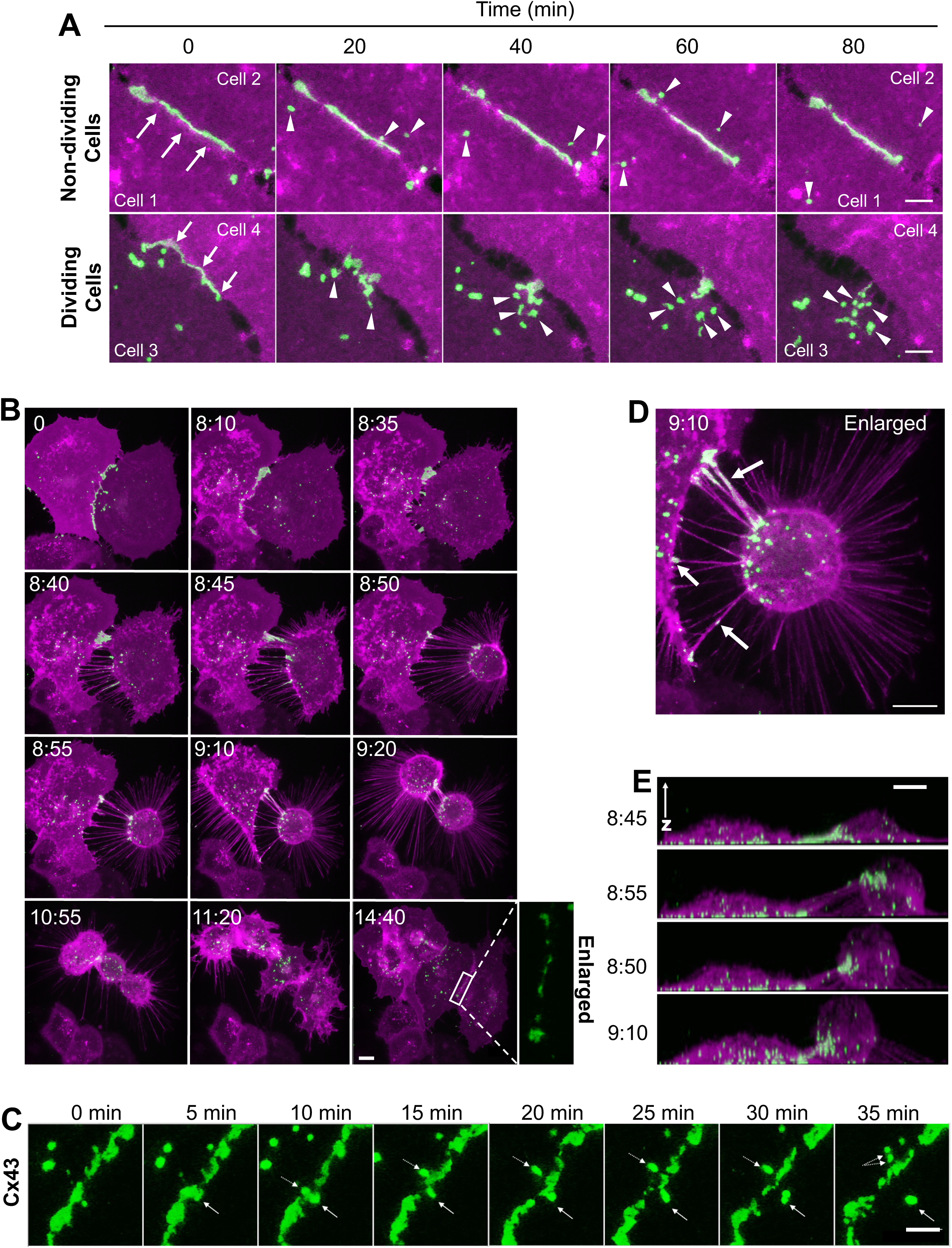
Live-cell confocal imaging of Cx43 dynamics throughout the cell cycle. (A) Time-lapse of gap junction plaque turnover in dividing and non-dividing cells. Gap junction plaques were monitored between pairs of interphase cells that later underwent mitosis (Dividing Cells) and pairs of interphase cells that did not divide over ≥ 16 hours of imaging (Nondividing Cells). Large plaques were present at cell-cell contacts at t=0 in both cell pair types (arrows). However, plaque internalization occurred more frequently, resulting in more annular gap junctions in the cytoplasm (arrowheads) of dividing cells than in non-dividing cells. In the montage shown, new annular gap junctions were similarly distributed between the cytoplasm of both cells (Cells 1 and 2). However, among the dividing cells, annular gap junctions were predominantly released into the cell that initially underwent mitosis, compared with the cell that divided later (Cell 4). After 80 minutes, little to no gap junction plaques remained at the surface of the dividing cells, whereas plaques between non-dividing cells showed no apparent change in size. (B) Time-lapse of gap junction dynamics as cells enter, undergo, and exit mitosis. Cells initially in interphase (t = 0:00) were observed to change shape, from flat to round (t = 9:20) and then divide to form daughter cells (t = 10:55) that flattened as they entered interphase (t = 14:40). New gap junction plaques were observed between daughter cells (t=14:40; also enlarged and shown with green channel only). (C) Gap junction plaque internalization into interphase cells in B. The gap junction plaque area has been enlarged, and the mEmerald channel (Cx43) is shown only. The mEmerald channel (Cx43) is shown only at 5-minute intervals for the first 1:20. Solid arrows follow the release of one annular gap junction over time, while dashed arrows follow the release and fission of a second annular gap junction. (D) Mitotic cell nanotubes from an area in B (t = 9:10) that has been enlarged to show Cx43-positive structures (arrows) visible along the mitotic nanotube as well as at both ends of the nanotube. (E) Mitotic nanotube changes shown with three-dimensional rotation of the images in A (t = 8:45 - 9:10). Note the change in Cx43-positive structure distribution within nanotubes over time and that the mitotic nanotubes remained above the substrate. Cells shown were synchronized, transfected to express mEmerald-Cx43 (green) and mCherry-CAAX (magenta), and imaged at 5-minute intervals. Scale bars = 5µm (A, C, D); 10µm (B, E).

To assess the dynamics and morphological changes of Cx43 from mitotic entry through exit, we characterized cell morphology throughout the cell cycle to reliably identify cell cycle stages using live-cell imaging, particularly when nuclear staining was not feasible for long-term imaging experiments (Figure S1). Cells in interphase were notably flat but rounded during metaphase, then underwent cytokinesis to divide into new daughter cells that eventually flattened (Figure S1A-D). These shape changes paralleled XY-dimensional cell area (Figure S1E, E’, F). Therefore, we used cell area and nuclear staining in some protocols to determine and correlate changes in gap junction structure with these cell cycle stages.

To assess changes in Cx43 distribution during mitosis, live imaging was performed on synchronized cells expressing mEmerald-Cx43 and mCherry-CAAX. At the start of imaging, cells were initially in interphase, and Cx43 was primarily localized to gap junction plaques on the plasma membrane, with smaller amounts present in cytoplasmic annular gap junction vesicles (Figure 2B, Supplementary Video 1). In the hours preceding mitosis, gap junction plaques were observed to decrease (Figure 2B), and this was largely driven by frequent gap junction internalization events that resulted in annular gap junction vesicle release into the cytoplasm (Figure 2C). Interestingly, in addition to the redistribution of Cx43 from gap junction plaques on the cell surface to annular cytoplasmic vesicles, plaques that remained on the cell surface during early cell rounding were observed to reposition into mitotic nanotubes (Figure 2B, D). The mitotic nanotubes formed bridge-like structures between the associated cells and remained above the substrate (Figure 2E). In addition, they appeared to form as a result of mitotic cells rounding and retracting from surrounding cells (Figure 2B). Once formed, mitotic nanotubes elongated as the mitotic cells progressed through mitosis, then gradually shortened and disappeared as the daughter cells flattened (Figure 2B, E). As the mitotic nanotubes lengthened and narrowed, Cx43-positive structures were observed at various locations along the lengths of the mitotic nanotubes (Figure 2B, D). After spreading, new plaques were observed to form between the daughter cells (Figure 2B; enlarged). Based on these imaging data, it is shown that mitotic nanotubes and associated Cx43 structures are dynamic and constantly reorganized throughout mitosis. However, enhanced imaging is necessary to more precisely track the movement of Cx43 structures associated with mitotic nanotubes over time.

### Movement of Gap Junction Structures within Mitotic Nanotubes

To further monitor dynamic changes in thin mitotic nanotubes, cells were imaged with a smaller Z-step size (0.25µm) to enable more detailed tracking of gap junction structures along the thin mitotic nanotubes. As the mitotic nanotubes lengthened, linear, gap junction plaque-like structures also lengthened to conform to the shape of these membrane extensions (Figure 3A). Although these Cx43-containing gap junction-like structures were linear, similar to the morphology of typical gap junction plaques, they were not located on the plasma membrane between two contacting cells and thus not positioned to allow for cell–cell communication. These linear Cx43 structures were usually observed at the early stages of cell rounding, and as cells continued to round and retract, Cx43 structures gradually fragmented into smaller structures (Figure 3A). Tracking the Cx43 structures showed that they moved in a unidirectional manner within the mitotic nanotube, but they did not seem to preferentially move toward or away from mitotic cells (Figure 3). In some cells, Cx43 structures were observed moving along mitotic nanotubes and transferring into the cytoplasm of one or both associated cells (Figure 3B). As seen in Figure 3B, a Cx43 structure at one end of a mitotic nanotube fragmented to form one annular gap junction-like structure that moved along the nanotube and entered the cytoplasm of the adjacent cell, while the residual Cx43 structure remained at one end of the nanotube, where it was positioned to potentially facilitate communication. The shape of the Cx43 structures did not appear to influence their movement across the mitotic nanotube, as both linear (Figure 3C) and blob-like (Figure 3B) structures were observed to move along these mitotic nanotubes.

**Figure 3.**
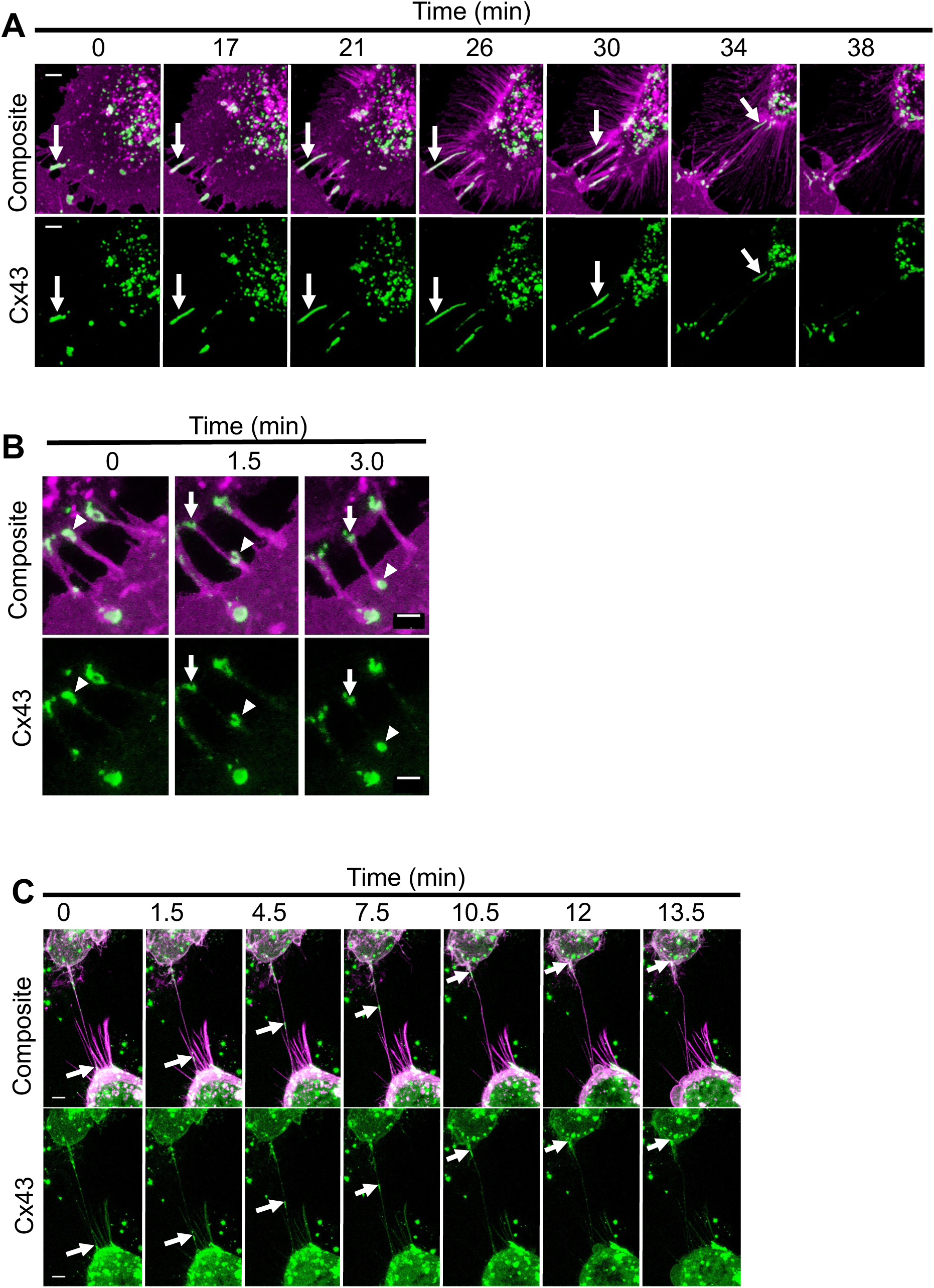
Confocal live-cell imaging of Cx43 structure dynamics within mitotic nanotubes. (A) Mitotic nanotubes in the area between a rounding mitotic cell and a neighboring cell. The solid arrow (0-34 min) represents a linear Cx43-positive structure within the mitotic nanotube. With time, the Cx43-positive structure elongates (0-26 min), fragments (30 min), and moves toward the dividing cell (34 min) before being lost from view (38 min). (B) Transfer of a Cx43-positive structure via mitotic nanotubes. A Cx43-positive structure (arrowhead), which resembles an annular gap junction, moved between two cells and changed shape within the mitotic nanotube before entering the cytoplasm. Some Cx43-positive material remained at one end of the mitotic nanotube (arrows). (C) Movement of a Cx43-positive structure along a mitotic nanotube. A mitotic-nanotube-associated Cx43 structure (arrows), initially at one end of the mitotic nanotube (0 min), moves along a straight path toward the other cell (1.5-10.5 min) and eventually enters the cytoplasm of the other cell (13.5 min). In this figure, cells were transfected to express mEmerald-Cx43 (green) and mCherry-CAAX (magenta) and confocal Z-stacks (step size = 0.25µm) were captured at 4-minute (A) or 1.5-minute intervals (B, C). Scale bars = 5µm (A, C); 2.5µm (B).

Based on these observations, Cx43-containing components moved within the nanotubes as well as between cells. In addition to cell-cell transfer of these Cx43-containing structures, Cx43-positive material was also observed at the ends of mitotic nanotubes, where they may be capable of facilitating gap junction-mediated cell–cell transfer through these extensions. Therefore, functional studies are necessary to determine whether mitotic nanotubes provide a mechanism for intercellular communication during mitosis despite the reduction in gap junction plaques.

### Quantitative Analysis of Cx43 Gap Junction Protein Distribution Changes Throughout the Cell Cycle

While it was also crucial to monitor gap junction trafficking events in individual cells, quantification of gap junction structure distribution was necessary to uncover trends in gap junction changes at the cell population level (Figure 4). Indeed, gap junction trafficking quantified by microscopy paralleled our findings from live-cell imaging.

**Figure 4.**
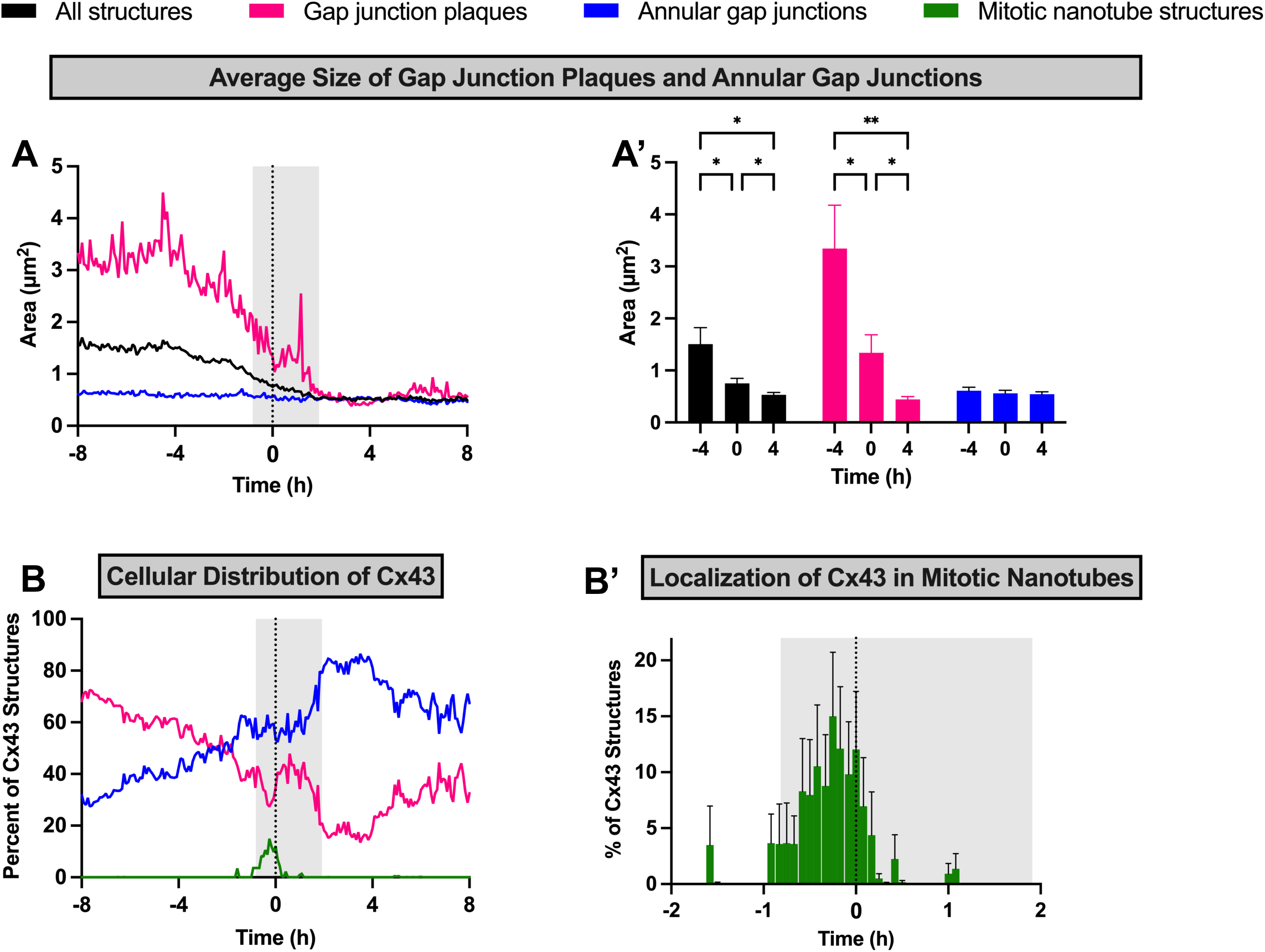
Quantitative analysis of gap junction structures throughout the cell cycle. For individual cells, each Cx43-positive structure was segmented, classified (as either a gap junction plaque, an annular gap junction, or associated with mitotic nanotubes), measured, and graphed. (A and A’) Average size of gap junction structures (B and B’). Cellular distribution of Cx43. Distribution was calculated as the total area of Cx43 structures in each compartment (gap junction plaques, annular gap junctions, or mitotic nanotubes) divided by the total area of Cx43 structures in the entire cell. In B’, axes in B are expanded to highlight changes in Cx43 localization in mitotic nanotubes. Graphs in A and B represent mean values versus time relative to metaphase (t =0). Graphs in A’ and B’ represent mean ± SEM at selected time points. Shaded gray area (A, B, B’) represents the average window during which cells were in mitosis (from initial rounding to cytokinesis). Asterisks represent the appropriate p-values: ≤0.033 (*), <0.002 (**), n = 20.

Changes in gap junction structure, size, and distribution were measured and expressed with respect to time relative to metaphase (Figure 4). In comparison to the average gap junction plaque size observed 4h before metaphase, the average gap junction plaque size was decreased by 60% at metaphase and 87% 4h after metaphase (Figure 4A, A’). The decrease in plaque size observed 4h post-metaphase coincided with the spreading of newly formed daughter cells and was significantly smaller than the plaque size observed between mother cells. These decreases in plaque size parallel observations from live-cell imaging of frequent gap junction internalization events (Figure 4A). This decrease in average plaque size four hours after metaphase could reflect downregulation of gap junctions due to internalization as well as the formation of new, relatively smaller gap junction plaques that aggregate to form into larger plaques, similar to those observed in the mother cells, over time (Figure 4A). Plaque internalization events were not observed as daughter cells flattened and formed new junctions. While the size of gap junction plaques decreased, the average size of annular gap junctions remained relatively constant throughout the course of the time-lapse analysis (Figure 4A, A’). This could indicate that annular gap junction processing events, like fusion and fission, that affect annular gap junction sizes are not significantly altered during mitosis.

To assess the changes in relative cellular distribution of Cx43, the percent of Cx43 in each compartment (i.e., gap junction plaques and annular gap junctions) relative to the whole cell was calculated (Figure 4B, B’). The percentage of Cx43 in gap junction plaques gradually decreased from 68% at 8 hours before metaphase to 48% 2h before metaphase, followed by a sharper decrease to 35% at metaphase. The percentage Cx43 in gap junction plaques decreased to a minimum of roughly 16% at 2h post-metaphase. These decreases in Cx43 localization to plaques are consistent with increased gap junction plaque internalization. Between 4h and 8h after metaphase, the percentage of Cx43 in gap junction plaques increased to 33%, consistent with increased plaque assembly in daughter cells (Figure 4B).

Interestingly, in addition to gap junction plaques and annular gap junctions, Cx43 was also temporarily redistributed to mitotic nanotubes (Figure 4B, B’). Localization of Cx43 to mitotic nanotubes was observed from 1.6h pre-metaphase to 1.2h post-metaphase, with maximal values observed 0.25h pre-metaphase when 15% of Cx43 localized to this compartment (Figure 4B’). This increase in Cx43 localization to the mitotic nanotubes corresponded to decreased localization to gap junction plaques, which is consistent with direct trafficking of Cx43 from plaques to mitotic nanotubes (Figure 4C). Overall, based on these quantitative analyses, it is speculated that throughout the cell cycle, alterations in various gap junction processing events occur, thereby influencing gap junction turnover and intercellular communication.

### The Role of Mitotic Nanotubes in Intercellular Communication

To assess the role of mitotic nanotubes in cell–cell communication, FRAP analysis was used (Figure 5). Rapid communication of calcein, a channel-permeable molecule, was observed between interphase cells that were in contact with one another, while isolated (non-contacting) cells failed to recover (Figure 5A). Isolated interphase cells were used as a negative control for cell–cell communication. Calcein recovery was observed in mitotic cells that were only in contact with neighboring cells via mitotic nanotubes. Calcein recovery rates for mitotic cells (8% ± 2.0) were significantly lower than rates for contacting interphase cells (34% ± 4.3) but higher than rates observed for isolated cells (1% ± 0.25) (Figure 5B, C). However, there were no differences in the mobile fractions (percent of unbleached molecules capable of moving into the bleached cell) for mitotic cells compared to contacting interphase cells (Figure 5D). Thus, the availability of calcein to move is similar; however, the rate of movement is slower than that observed between contacting interphase cells.

**Figure 5.**
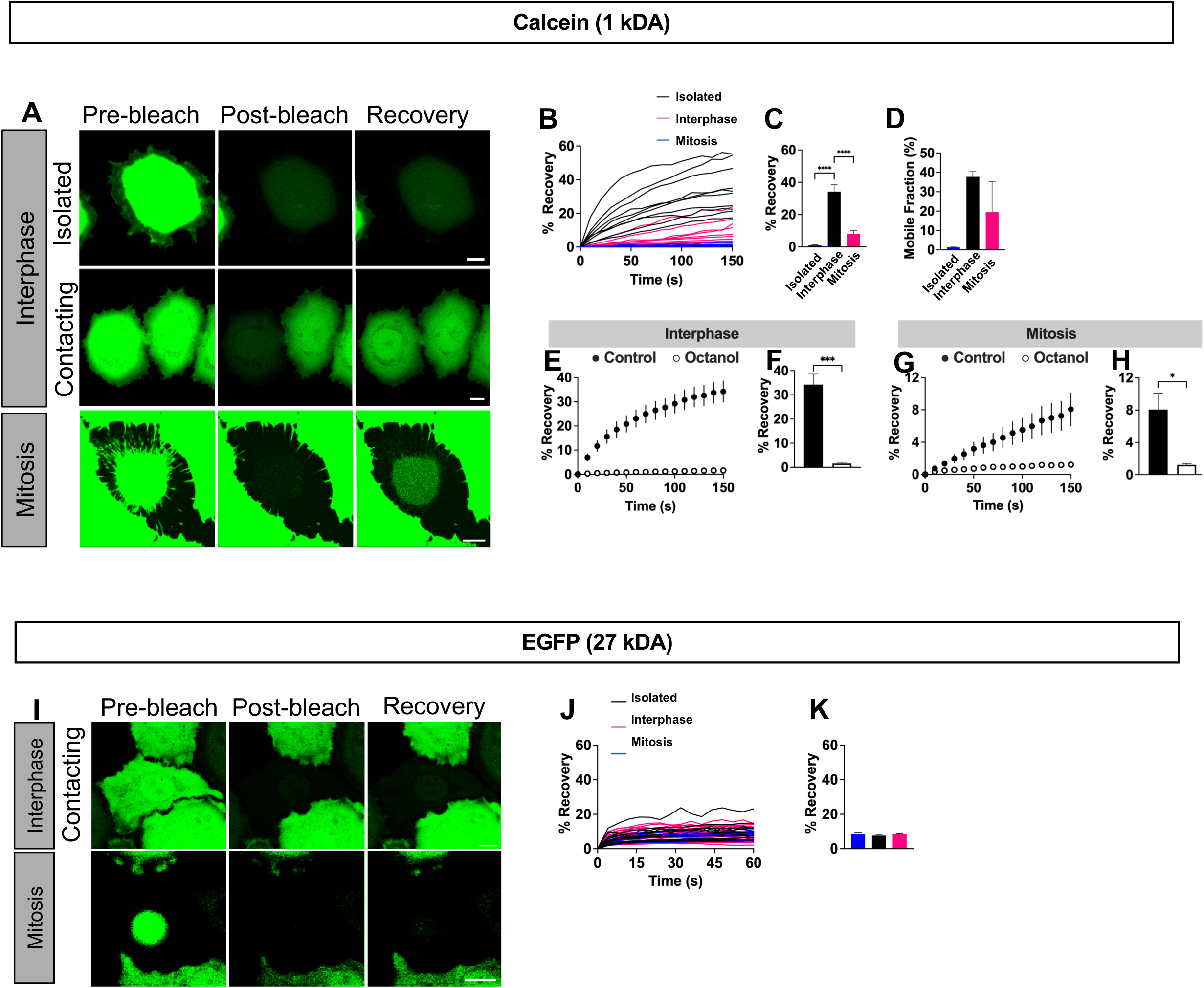
Fluorescence recovery after photobleaching (FRAP) analysis of intercellular communication of calcein and EGFP in cells in interphase and mitosis. (A) Confocal images of calcein-loaded cells before, immediately after and 150s after bleaching (recovery). (B-D) Calcein communication in untreated cells. (E-H) Calcein communication in octanol-treated and untreated cells. (I) Confocal images of EGFP-expressing cells before, immediately after and 60s after bleaching (recovery). (J, K) Communication of EGFP in untreated cells. Each line in B and J represents a single cell; graphs in E and G represent mean fluorescence recovery ± S.E.M over time; graphs in C,F,H,K) represent fluorescence at the end of recovery; D represents the mobile fraction (Ymax) after fitting to a nonlinear recovery curve. Asterisks (*) in represent the appropriate p-values: ≤0.033 (*), <0.001 (***), <0.0001 (****); n ≥ 3 cells per group. Scale bars (A, M) = 10µm.

To determine the role of gap junction-mediated communication in mitotic nanotube-mediated calcein recovery, gap junction channel function was inhibited, and communication was measured with FRAP analysis. Recovery rates were decreased between contacting interphase cells and between mitotic cells in contact with surrounding cells via mitotic nanotubes following treatment with octanol, which blocks gap junction channels (Figure 5E-H). Thus, mitotic nanotube-mediated cell–cell communication of calcein was inhibited when gap junction channels were compromised, which suggests that mitotic nanotubes allow for the movement of small molecules in a gap-junction-dependent mechanism.

To determine if mitotic nanotubes could allow the passage of molecules larger than what can move through a gap junction channel pore (1.2kDA), we assessed intercellular communication of EGFP (Figure 5M-O). At 27kDA, EGFP is too large to pass through gap junction pores. In interphase cells, no recovery was observed after bleaching. On average, mitotic cells did not recover significant EGFP fluorescence above background following photobleaching (Figure 5M-O). Thus, mitotic nanotubes facilitate intercellular communication via gap junctions; when gap junction function was inhibited, communication was reduced. However, mitotic nanotubes do not appear to allow the transfer of molecules, such as EGFP, that are too large to pass through gap junction channels.

### Role of Cx43 in Mitotic Nanotube Formation and Function

To investigate the role of Cx43 in mitotic nanotube formation and function, cells were transfected with Cx43 siRNA to knock down expression, or with non-targeted control siRNA, and then assessed with Western blotting and immunocytochemistry to validate knockdown efficiency (Figure 6A, B). Mitotic nanotubes were often absent in Cx43 knockdown cell populations (Figure 6C, D). In contrast, cells transfected to express the control siRNA formed mitotic nanotubes (Figure 6C, D) that were similar in appearance to those observed in non-transfected controls (Figure 1F). In Cx43 knockdown populations, mitotic cells rounded in preparation for division but, in many cases, did not retract from neighboring cells and instead remained in close contact (Figure 6C, D). To better characterize the dynamic changes resulting from reduced Cx43 expression, cells co-transfected with either control or Cx43 siRNA and the membrane marker mCherry-CAAX were imaged at 5-minute intervals (Figure 6E). Mitotic nanotubes were observed to form and elongate as control cells rounded during mitosis (Figure 6E). However, in Cx43 knockdown cell populations, cells frequently progressed through the entire mitotic process without retraction from adjacent cells or mitotic nanotube formation (Figure 6E) while others were seen to contract and form nanotubes (not shown). We identified cells in metaphase that formed mitotic nanotubes in Cx43 knockdown and control cell populations, and measured calcein cell–cell communication (Figure 6 H,I). In knockdown cells, dye communication was reduced by 65% in interphase cells and by 80% in mitotic cells relative to control cells at the corresponding cell cycle stages (Figure 6F-I). These results are consistent with Cx43 playing pivotal role in both mitotic nanotube formation and cell–cell communication rates.

**Figure 6.**
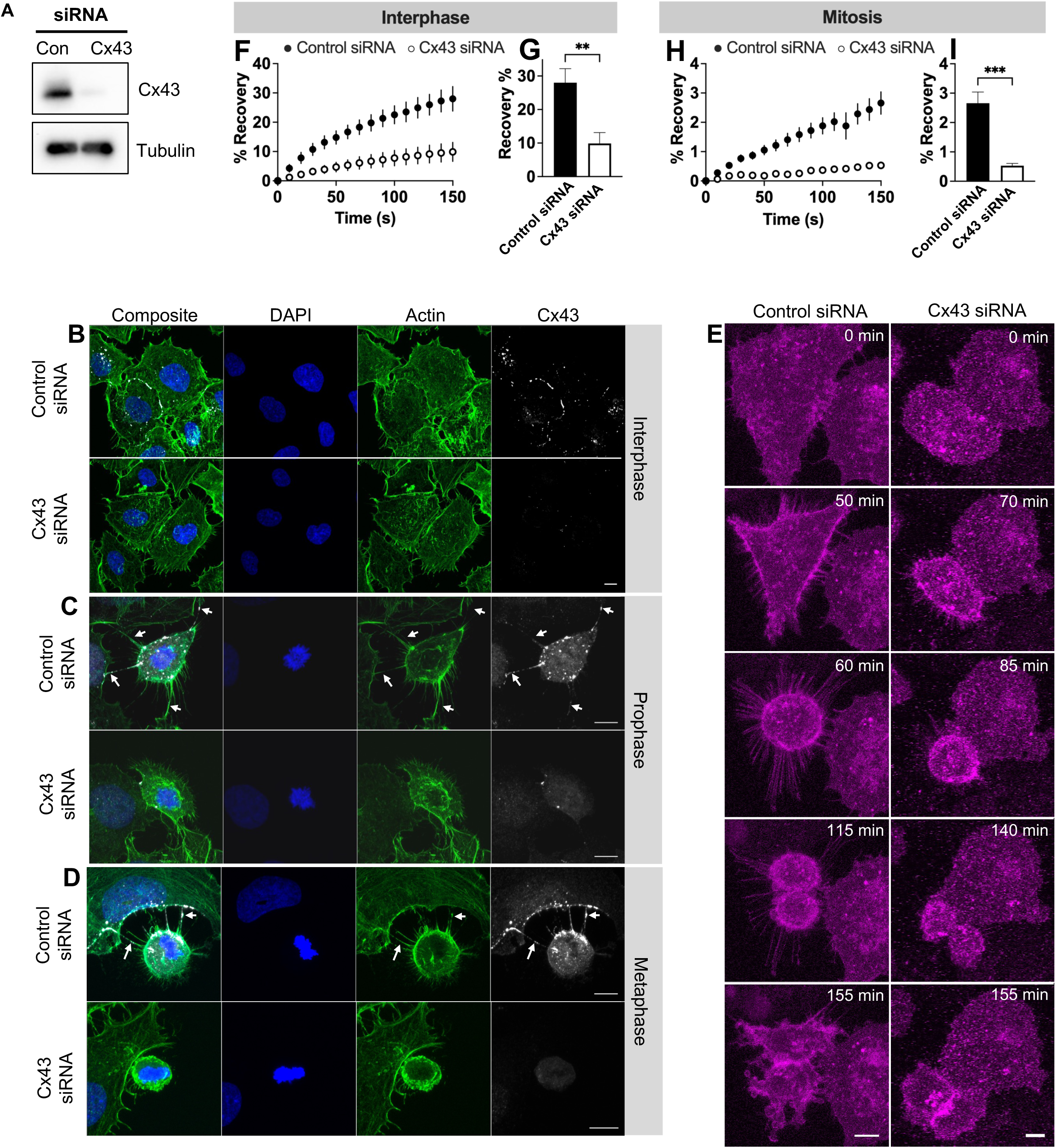
Effect of Cx43 knockdown on mitotic nanotube structure and function. (A) Western blot of cell lysates after transfection with control siRNA or Cx43 siRNA. (B-D) Immunocytochemistry of control and knockdown cells during interphase (B), prophase (C) and metaphase (D). Control cells formed typical mitotic nanotubes (arrows) during mitosis but Cx43 knockdown cells were observed without mitotic nanotubes and mitotic cell bodies remained in close proximity or attached to adjacent cells. (E) Live-cell images of mitosis. Cells were synchronized with a double-thymidine block, transfected with mCherry-CAAX, and control or Cx43 siRNA then imaged at 5-minute intervals. Cells initially in interphase, changed shape and divided to form daughter cells with time. Mitotic nanotubes were typically observed to form in control cells but formed less frequently in knockdown cells as cells remained in contact with adjacent cells during mitosis. Scale bars = 10µm. (F-I) Cell-cell communication rates in control and knockdown cells. Calcein communication was measured with FRAP. In Cx43 knockdown cells, communication was reduced by 65% during interphase and 80% during mitosis, compared to control cells at the same stage. Graphs in F and H represent fluorescence recovery over time; graphs in G and I represent fluorescence recovery 150s after bleaching. Data = mean ± S.E.M. Asterisks = p: <0.002 (**), <0.001(***); n ≥ 6 cells per group.

### Regulation of Mitotic Nanotubes Cell-Cell Communication by Rho Kinase (ROCK)

Given that mitotic nanotubes are actin-based structures, studies were performed to elucidate the role of F-actin reorganization. To investigate this, cells were treated with the ROCK inhibitor Y-27632, which disrupts actin filaments [27] and impairs mitotic cell rounding [28]. Gap junction plaques were consistently larger in the ROCK inhibitor treated cells compared to controls (Figure 7A). Furthermore, in control cells, mitotic cell rounding was observed early in prophase, with more pronounced rounding in metaphase (Figure 7B, C). However, in cells treated with Y-27632, the degree of cell rounding was reduced such that the cells appeared flatter and more spread during prophase and metaphase (Figure 6B, C). Further, while control cells exhibited the typical elongated mitotic nanotube structures, Y-27632-treated cells formed comparatively shortened mitotic nanotubes, and their cell bodies remained in relatively a closer contact with adjacent cells (Figure 7B, B’, C, E). While control and inhibitor-treated cells exhibited similar numbers of mitotic nanotubes during prophase, Y-27632-treated metaphase cells exhibited significantly more mitotic nanotubes than untreated metaphase controls (Figure 7D). Taken together, these results suggest that ROCK is required for mitotic cell rounding, and that when ROCK is inhibited, mitotic nanotubes form initially, but these structures are shorter than those in DMSO-treated controls. ROCK inhibition is therefore associated with lack of nanotube elongation and with prolonged persistence of mitotic nanotubes as cells progress to metaphase. Despite differences in mitotic nanotube morphology, Cx43 was associated with the mitotic nanotubes in both control cells and Y-27632-treated cells (Figure 7C, E). To evaluate whether intercellular communication was altered in these cells following ROCK inhibition FRAP analysis was used.

**Figure 7.**
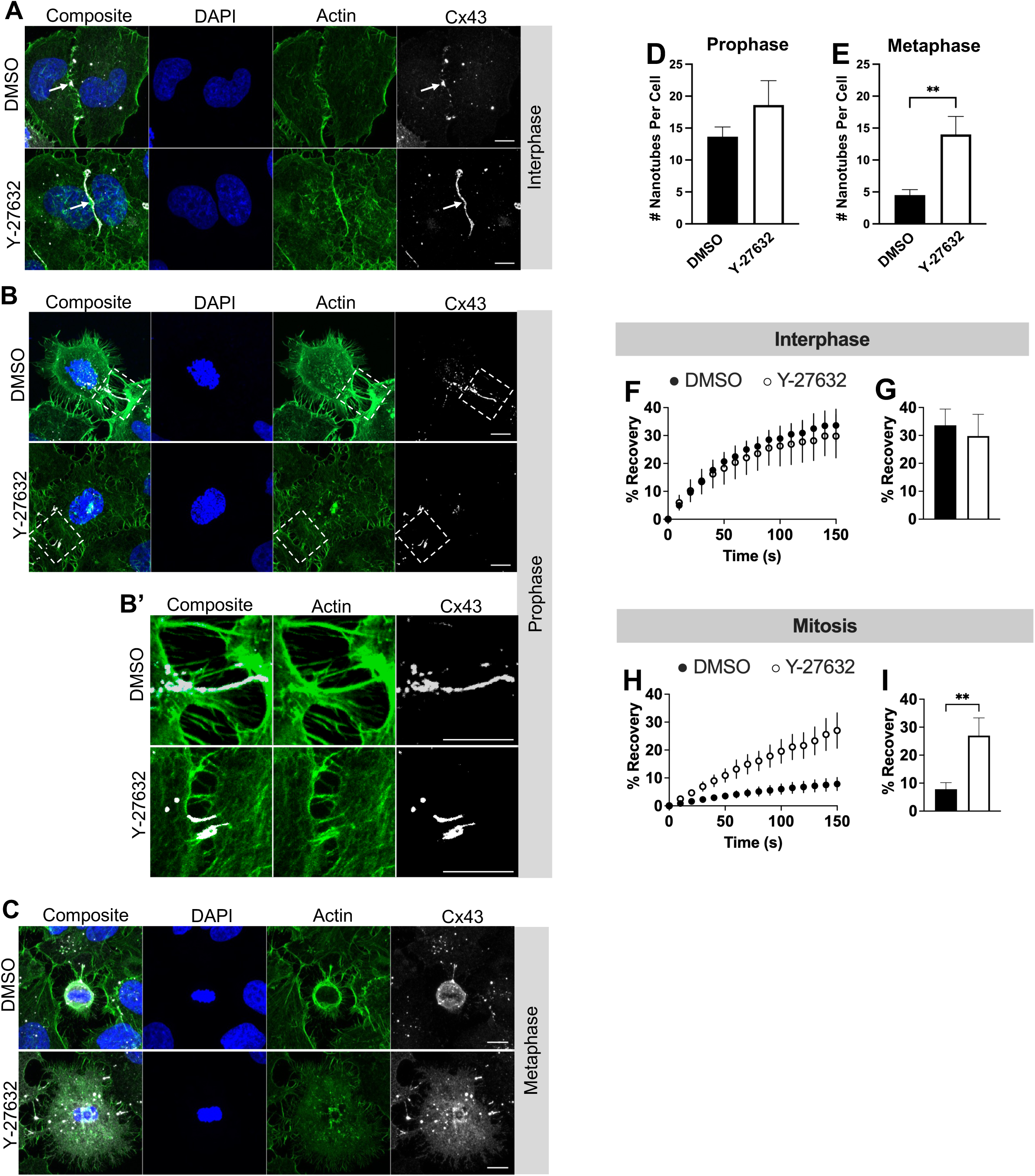
Effect of Rho kinase (ROCK) inhibition on mitotic nanotube structure and function. (A - C) Immunocytochemistry of SW-13 cells treated with DMSO (control) and the ROCK inhibitor Y-27632. During interphase, larger gap junction plaques were observed (arrows in A). Mitotic cell rounding was observed early in prophase, with additional rounding in metaphase in DMSO-treated cells. However, Y-27632-treated cells maintained a well-spread morphology during prophase and metaphase (B, C). In addition, mitotic nanotubes in Y-27632 cells appeared shorter than those in control cells (B’). Scale bars = 10µm. (D,E) Graphs of the number of mitotic nanotubes per cell. (F-I) Analysis of calcein communication in DMSO- and Y-27632-treated cells. Graphs in F and H represent fluorescence recovery over time; graphs in G and I represent fluorescence recovery 150s after bleaching. Data = mean ± S.E.M. Asterisks = p: <0.03 (*), <0.002 (**); n ≥ 3 cells per group.

Although Y-27632-treated interphase cells exhibited larger gap junction plaques than control cells (Figure 7A), calcein transfer rates were similar between control and inhibitor-treated interphase cells (Figure 7F, G). However, Y-27632-treated mitotic cells facilitated calcein communications at rates significantly higher than control mitotic cells. Therefore, these findings are consistent with ROCK inhibition preventing mitosis-associated reductions in gap junction-mediated communication.

## Discussion

In this study, we characterized the dynamics of gap junction protein trafficking during cell entry into, passage through, and exit from mitosis. In addition to demonstrating the behavior of gap junction plaques and annular gap junctions, we also characterize connexin trafficking to poorly characterized actin-based membrane extensions, termed mitotic nanotubes, that house Cx43 gap junction plaque material during mitosis, thus supporting a role for these membrane projections in preserving connexin complexes as cells undergo division. These nanotubes undergo dynamic remodeling during mitosis but are transient and disappear as mitosis ends. We also uncover a novel role for Cx43-containing mitotic nanotubes in facilitating gap junction intercellular communication, thereby bridging a critical gap in our understanding of how these membrane extensions contribute to intercellular communication even as gap junction plaques are removed from the cell. Although lengthening of the mitotic nanotubes is reduced when F-actin organization is perturbed by inhibition of ROCK, mitosis-associated reductions in cell-cell communication are prevented in the presence of this inhibitor, indicating that in addition to Cx43 function, the dynamic remodeling of mitotic nanotubes may also play a role in regulating cell-cell communication between mitotic cells and their neighbors. Overall, we reveal how changes in Cx43 trafficking and the remodeling of mitotic nanotubes work in tandem to regulate cell–cell communication during mitosis (Figure 8).

**Figure 8.**
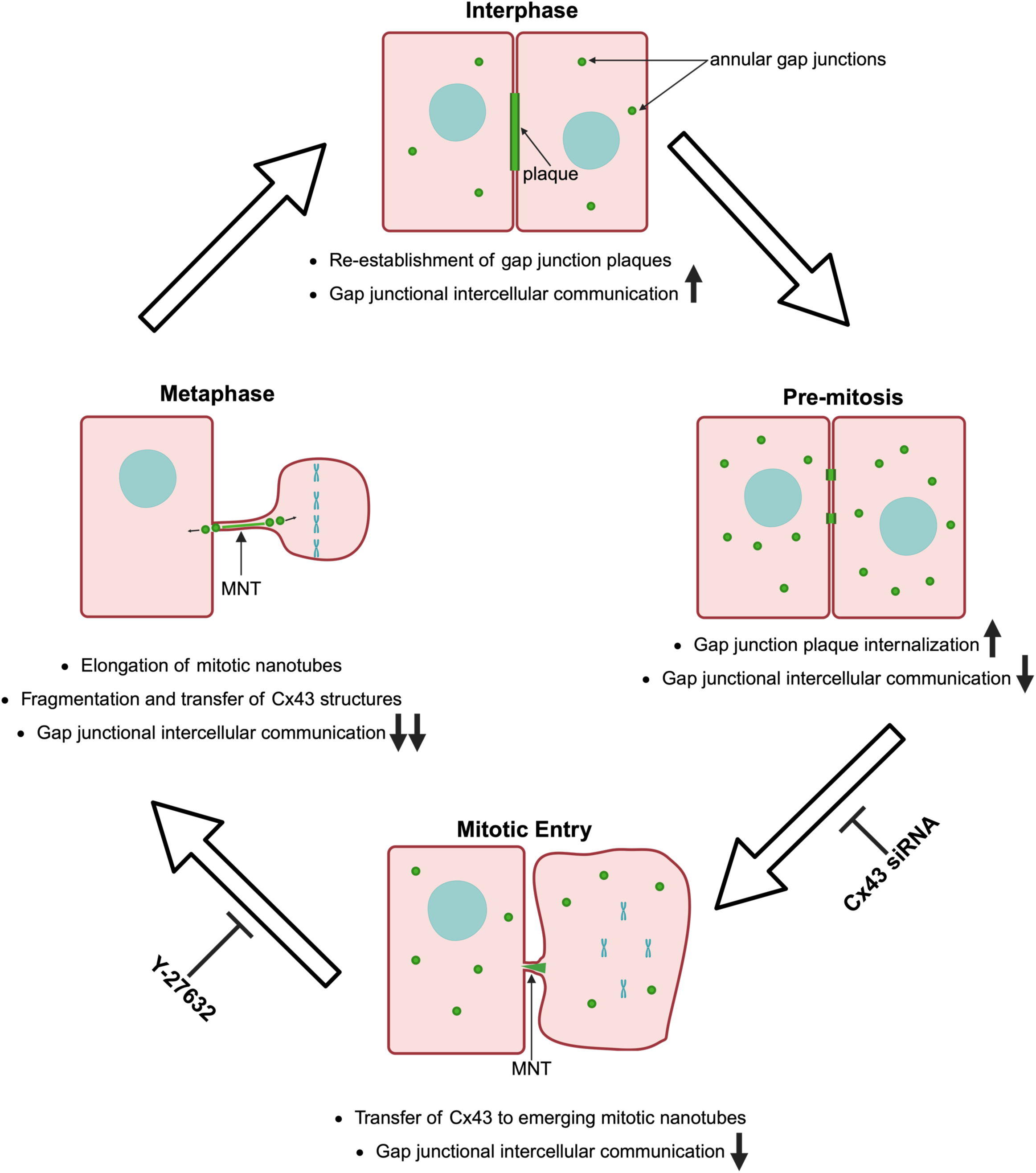
Graphical abstract illustrating alterations to Cx43 trafficking and their functional consequences on gap junction-dependent intercellular communication as cells prepare for, undergo and exit mitosis. During interphase, cell-cell communication occurs rapidly via large gap junction plaques. In cells that are still in interphase but preparing for division (Pre-Mitosis), gap junction plaque internalization is upregulated, resulting in increased annular gap junction release and decreased intercellular communication. As cells enter mitosis, plaques that remain on the cell surface are transferred to emerging mitotic nanotubes. The formation of these nanotubes is reduced when cells are transfected with Cx43 siRNA. Cells progressing further through mitosis (around Metaphase) continue to undergo mitotic cell rounding and mitotic nanotubes elongate with time. The Cx43 structures associated with the nanotubes are fragmented and transferred to the cytoplasm of either cell as annular gap junction-like structures or deposited at the ends of the mitotic nanotubes. Gap junction-dependent intercellular communication occurs via mitotic nanotubes but is rates are reduced compared to rates between interphase cells. However, treatment with the ROCK inhibitor Y-27632 prevents lengthening of MNTs, mitotic cell rounding and reductions in cell-cell communication during mitosis, suggesting a role for ROCK and cytoskeletal remodeling in regulating mitotic nanotube function. As cells exit mitosis, gap junction plaques assemble in the newly formed daughter cells and cell-cell communication rates increase. Green structures represent Cx43-positive structures. MNT = mitotic nanotube. This figure was created in BioRender: Cooper, A. (2026) https://BioRender.com/yphb61r

The dynamic reorganization of gap junction plaques is crucial for mediating cell–cell communication. Despite reductions in gap junction communication during mitosis [17, 18], there may be a need for some degree of communication during mitotic cell reorganization. Although tight junctions have been shown to remain intact during mitosis [29–31], previous studies have demonstrated the disassembly of adherens junctions during mitotic cell rounding and have suggested the molecular basis and necessity of this disassembly. While studies document the loss of plaques during mitosis [19, 21] or reduced cell coupling during mitosis [17, 18], visualization of these dynamic changes in Cx43 localization as cells prepare for, undergo, and complete mitosis remains limited. Using live-cell confocal imaging we achieved three-dimensional visualization of cells undergoing robust shape changes, with high spatial and temporal resolution, thereby enabling reliable quantification of the dynamics of Cx43-positive structures across the cell cycle. We demonstrate that in addition to gap junction plaque internalization processes that result in annular gap junction release into the cytoplasm, gap junction plaque-derived Cx43 is also sequestered within the transient mitotic nanotubes. In many cases, the gap junction components within mitotic nanotubes appear to be delivered to one or both contacting cells after mitosis. Additionally, as cells exit mitosis and form new daughter cells, plaques are frequently reestablished in areas where gap junction plaques were observed prior to mitosis, and therefore near regions where Cx43 structures were observed at the ends of mitotic nanotubes between the dividing cells and their neighbors. This could suggest that these portions of the plasma membrane are targeted areas, or hotspots, for gap junction plaque assembly. We also suggest, based on these findings, that Cx43 associated with mitotic nanotubes might participate in the rapid restoration of gap junction plaques at the plasma membrane after mitosis in a mechanism that involves recycling of Cx43 rather than *de novo* Cx43 synthesis and gap junction assembly.

Morphologically, mitotic nanotubes resemble tunneling nanotubes, which have been shown to allow the transfer of organelles, viruses, and signaling molecules between cells [32–37]. Like tunneling nanotubes, the mitotic nanotubes were observed to have a tubular morphology and to bridge two cells. In the case of mitotic nanotubes, these extensions bridge mitotic cells to cells with which they were in contact before the onset of division. In contrast, tunneling nanotubes have been reported to form between cells that are often several cells apart and have been described mainly in interphase cells, not during mitosis [34, 38]. The tunneling nanotube has been demonstrated to typically contain a single Cx43 structure at one end of the extension [39, 40] rather than multiple Cx43 structures at various points along its length as seen in mitotic nanotubes. In addition, Cx43 plays a role in the intercellular transfer of small materials and mitochondria via tunneling nanotubes [39, 41–44]. Within mitotic nanotubes however, gap junction structures conformed to the shape of the extensions as they emerged. With time, the Cx43 structures fragmented and were dispersed to various sites along the extensions eventually remaining at the ends of the nanotube or being transferred to the cytoplasm of the adjacent cell. The Cx43-positive structures that were transferred between cells were similar in size, shape, and appearance to annular gap junctions suggesting that gap junction remodeling within nanotubes shares mechanistic features with the formation of annular gap junctions during plaque internalization.

The finding that mitotic nanotubes form bridges between dividing cells and their neighbors is consistent with a potential role for these structures in preserving gap junction intercellular communication during mitosis. Based on our functional FRAP studies, we demonstrate that cells can communicate small, gap-junction-permeable molecules via mitotic nanotubes. Still, communication rates are slower than those observed via gap junctions between interphase cells. Since initial calcein assays performed did not allow us to distinguish whether communication via mitotic nanotubes was dependent on gap junction channels, additional assays were conducted to measure calcein communication in the presence of known gap junction inhibitors and to assess communication of EGFP, which is a larger, non-gap junction channel-permeable protein. Based on these studies, we confirm that the mitotic nanotubes facilitate communication in a gap junction-dependent manner. While cell–cell communication of EGFP was not observed via mitotic nanotubes in our experiments, the ability of mitotic nanotubes to facilitate communication of larger molecules or organelles in other contexts cannot be completely ruled out. Further, it is possible that during mitosis mitotic nanotubes can transition between open and closed states due to fusion between the mitotic nanotube membrane and the plasma membrane of the contacting cell.

Interestingly, downregulation of ROCK and Cx43 have opposing roles on mitotic nanotube function which suggests that they both regulate the function of mitotic nanotubes but in different mechanisms. Knockdown of Cx43 had an inhibitory effect on the formation of mitotic nanotubes and when nanotubes were present, nanotube communication rates were significantly reduced compared to rates in control cells. In Cx43 knockdown populations, reduced communication via mitotic nanotubes is consistent with Cx43 forming functional gap junctions at the ends of mitotic nanotubes. Based on imaging data, it could be suggested that Cx43 regulates the formation of the mitotic nanotubes. It is also possible that the plaques that remain on the cell surface during cell rounding mark key sites for mitotic nanotube formation and therefore when Cx43 expression is reduced, fewer nanotubes are formed.

Cytoskeletal organization via ROCK also regulates cell-cell communication via mitotic nanotubes. Based on inhibition data it is suggested that ROCK and its downstream effectors influence mitotic nanotube lengthening either by stabilizing actin filaments or by promoting mitotic cell retraction. However, the shortened nanotubes observed in inhibitor-treated cells allowed for more rapid communication of gap junction-permeable molecules than in control cells, which typically have lengthened mitotic nanotubes. Therefore, while we establish that mitotic nanotubes facilitate intercellular communication, perturbing the dynamic lengthening that occurs under normal conditions appears to enhance communication. Since ROCK inhibition does not affect the ability of cells to communicate during interphase, this change in communication is unlikely to be due to changes in gap junction channel gating. However, increased communication observed in mitotic cells during ROCK inhibition could be a result of the altered positioning or morphology of Cx43 structures associated with mitotic nanotubes. Specifically, partially elongated Cx43 components in ROCK-inhibited cells may allow for more efficient communication than the typical, lengthened Cx43 components in the mitotic nanotubes of control cells. This finding further highlights the need to visualize the 3D ultrastructure of Cx43 structures associated with mitotic nanotubes and suggests that these nanotubes are influenced by actin remodeling changes throughout the mitotic process.

The transient nature of mitotic nanotubes supports their role as an adaptive mechanism for mitotic cell–cell communication during mitotic cell remodeling rather than a permanent feature of intercellular communication. Their disappearance at the end of mitosis suggests that, unlike tunneling nanotubes, they serve a temporally restricted function, ensuring that cell–cell communication is retained and that the Cx43 components are appropriately redistributed upon cytokinesis. Future studies will be essential to identify the materials being communicated via mitotic nanotubes, characterize the ultrastructure of connexin structures associated with mitotic nanotubes, determine the roles of connexin phosphorylation and gap junction pore gating in this communication, and determine the impact of gap junction material being sequestered in these structures on mitosis and on post-mitotic gap junction reassembly.

In conclusion, our study uncovers previously unrecognized details of gap junction protein trafficking across the cell cycle and the role of gap junction protein-associated mitotic nanotubes in the regulation of intercellular communication during mitosis. These findings provide new insights into how cells preserve gap junction components during division and may have implications for understanding tissue homeostasis and mitotic regulation in both normal and pathological contexts.

## Acknowledgements

This work was funded by a National Science Foundation grant (#MCB2011577) awarded to S.A.M and UNCF/Bristol-Myers Squibb E.E. Just Faculty Fund, Career Award at the Scientific Interface (CASI Award) from the Burroughs Welcome Fund (BWF) ID # 1021868.01, BWF Ad-hoc Award, NIH Small Research Pilot Subaward 5R25HL106365-12 from the National Institutes of Health PRIDE Program, DK020593, Vanderbilt Diabetes and Research Training Center for DRTC Alzheimer’s Disease Pilot & Feasibility Program, CZI Science Diversity Leadership grant number 2022-253529 from the Chan Zuckerberg Initiative DAF, an advised fund of the Silicon Valley Community Foundation to A.H.J. We would like to thank Simon Watkins and University of Pittsburgh Center for Biologic Imaging for confocal microscopy equipment and technical support, and Filip Sluzewski, Julie Theriot and Matheus Viana at the Allen Institute for Cell Science and George Langford at the PAIR-UP Imaging Science Program for training in image analysis.

## Figure Legends

**Supplementary Figure 1.**
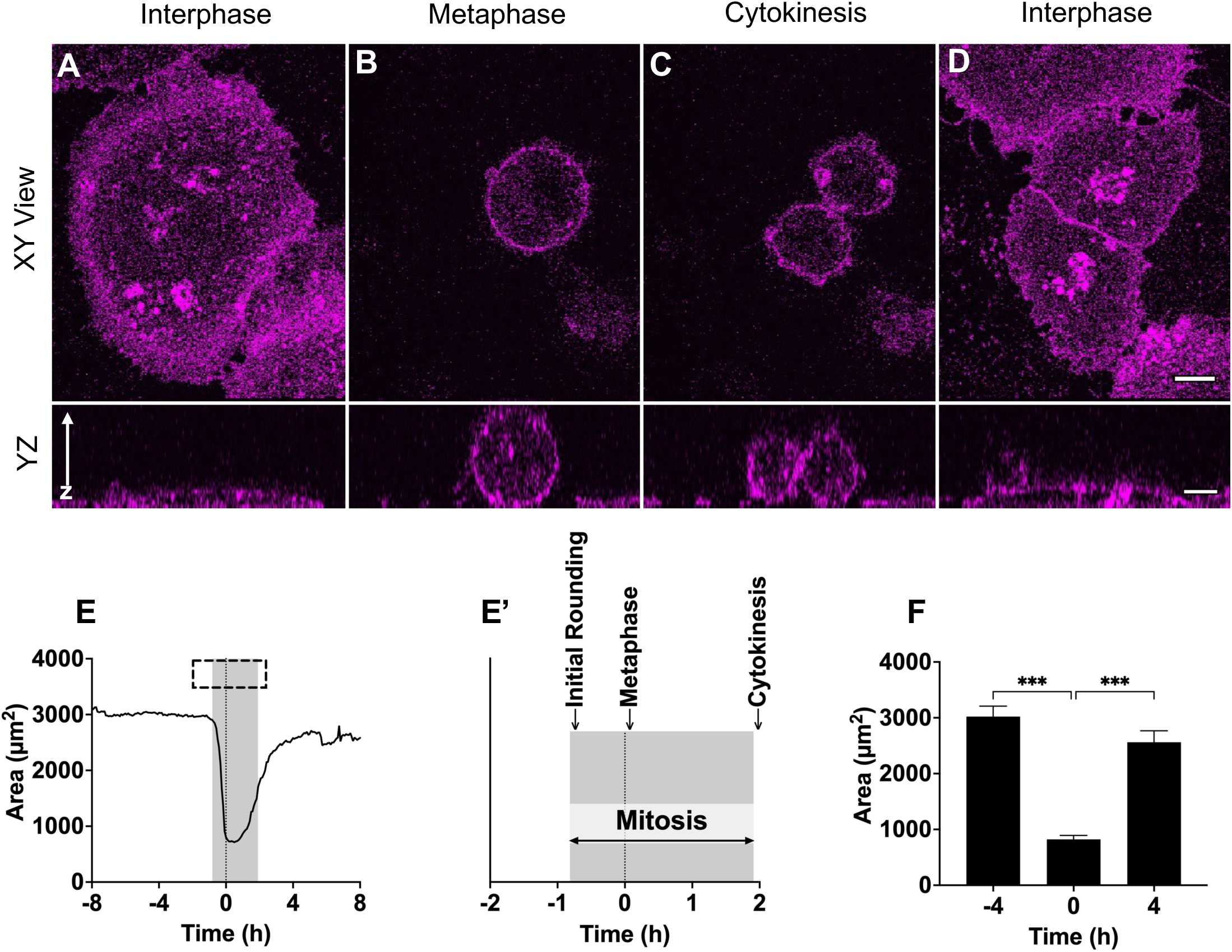
Analysis of cell shape changes during mitosis. (A-D) Confocal time-lapse images of a cell undergoing mitosis. Cells preparing to undergo division (A) were observed to round (B) and form daughter cells (C), which spread (D). Cells were synchronized and transfected to express the membrane marker mCherry-CAAX, then imaged at 5-minute intervals. Scale bars = 10µm. (E) Quantification of average cell area over time. Cell area decreased sharply as cells approached metaphase (t=0) and gradually returned to pre-mitotic levels thereafter. (E’) Enlargement of the x-axis in Y to show the average time during which mitotic events occurred. The gray area represents the average period that the cells are in mitosis, and the vertical line (t=0) represents metaphase. On average, initial mitotic cell rounding was observed 0.82 ± 0.33 hours before metaphase, and cytokinesis was observed 1.9 ± 0.36 hours after metaphase. (F) Graph of cell area at metaphase (t = 0), and at 4 hours before and after metaphase. Bar graph represents mean ± S.E.M.; n = 20. Times shown in E and F are relative to the estimated metaphase time point based on cell area. Asterisks (*) represent the appropriate p-values: 0.12 (ns), 0.033 (*), 0.002 (**), <0.001 (***).

